# An unusual MHC molecule generates protective CD8+ T cell responses to chronic infection

**DOI:** 10.1101/2020.02.03.932848

**Authors:** A. Tsitsiklis, D.J. Bangs, L.K. Lutes, S-W. Chan, K. Geiger, A.J. Modzelewski, L. Labarta-Bajo, Y. Wang, E.I. Zuniga, S. Dai, E.A. Robey

## Abstract

The CD8+ T cell response to the intracellular parasite *Toxoplasma gondii* varies dramatically between mouse strains, resulting in differences in control of the parasite. Protection in BALB/c mice can be attributed to an unusually strong and protective MHC-1 L^d^-restricted CD8+ T cell response directed against a peptide derived from the parasite antigen GRA6. The MHC-1 L^d^ molecule has limited peptide binding compared to conventional MHC molecules such as K^b^ or D^b^, which correlates with polymorphisms associated with “elite control” of HIV in humans. To investigate the link between the unusual MHC-1 molecule L^d^ and the generation of “elite controller” CD8+ T cell responses, we compared the GRA6-L^d^ specific T cell response to the well-studied OVA-K^b^ specific response, and demonstrated that GRA6-L^d^ specific T cells are significantly more protective and resistant to exhaustion in chronic *T. gondii* infection. To further investigate the connection between limited peptide presentation and robust T cell responses, we used CRISPR/Cas9 to generate mice with a point mutation (W97R) in the peptide-binding groove of L^d^ that results in broader peptide binding. We investigated the effect of this L^d^ W97R mutation on another robust L^d^-restricted response against the IE1 peptide during Murine Cytomegalovirus (MCMV) infection. This mutation leads to an increase in exhaustion markers in the IE1-L^d^ specific CD8+ T cell response. Our results indicate that limited peptide binding by MHC-1 L^d^ correlates with the development of robust and protective CD8+ T cell responses that may avoid exhaustion during chronic infection.

## Introduction

Individuals differ dramatically in their ability to mount CD8+ T cell responses against intracellular infections, as exemplified by Human Immunodeficiency Virus (HIV) infected individuals who can control the virus without anti-retroviral therapy (Deeks and Walker 2007). HIV “elite control” is strongly correlated with polymorphisms in genes encoding the major histocompatibility complex (MHC) class I proteins (Goulder and Walker 2012). While the ability of elite control-associated alleles, such as Human Leukocyte Antigen (HLA)-B57 and B27, to induce strong CD8+ responses undoubtedly depends on their ability to bind particular viral peptides, these MHC alleles are also associated with autoimmunity and resistance to other infections, suggesting that more general properties of these molecules may underlie their ability to generate potent CD8+ T cell responses (Bowness 2015; Hraber, Kuiken, and Yusim 2007; A. Y. Kim et al. 2011). Moreover, while potent CD8+ T cell responses are typically characterized by high affinity binding of T cell receptors (TCRs) for pathogen-derived peptides presented by MHC-1 proteins, other factors such as peptide affinity for MHC, T cell precursor frequency, and antigen density are also important (Tscharke et al. 2015; Jenkins and Moon 2015). Overall, the factors that determine the efficacy of CD8+ T cell responses, and why potent responses are linked to certain MHC-1 alleles, remains poorly understood.

Recent studies have revealed that the level of self-reactivity is another important factor in determining the strength of a T cell response. T cells undergo functional tuning in the thymus based on the strength of the self-pMHC signals they receive (Hogquist and Jameson 2014). CD5 is a key player in the tuning process, and also serves a robust marker of the degree of self-reactivity of naïve T cells (Azzam et al. 2001; Persaud et al. 2014; Fulton et al. 2015; Mandl et al. 2013). T cells with higher self-reactivity (CD5^high^) are generally superior in their responses to foreign antigens, with higher antigen-driven expansion, IL-2 production, and reactivity to cytokines compared to T cells with low self-reactivity (CD5^low^) (Weber et al. 2012; Mandl et al. 2013; Persaud et al. 2014; Fulton et al. 2015; Cho et al. 2016; Hogquist and Jameson 2014). In one study, a CD5^low^ CD4+ T cell clone displayed resistance to activation-induced cell death and superior expansion during secondary challenge (Weber et al. 2012). This suggests that high versus low self-reactivity T cells may be designed to fulfill distinct roles in different infection settings. However, the role of CD8+ T cells with low self-reactivity remains unknown.

While different MHC alleles vary in their ability to bind to particular antigenic peptides, there are indications that other properties of MHC-1 molecules may also play a role in shaping T cell responses. Human HLA-B alleles differ in several properties including their dependence on components of the antigen processing machinery, stability in absence of peptides, peptide loading efficiency, and the broadness of their specificity for peptides (Yarzabek et al. 2018; Geng, Zaitouna, and Raghavan 2018; Košmrlj et al. 2010; Chappell et al. 2015). It has been proposed that, while the majority of MHC-1 molecules are “generalists” able to bind a large set of peptides and produce adequate CD8+ responses to a large number of pathogens, certain MHC-1 molecules with distinctive peptide binding properties may serve as “specialists” that sacrifice broad coverage in exchange for the potential to make potent T cell responses to specific pathogens (Chappell et al. 2015; Košmrlj et al. 2010). Notably, most mouse model studies of CD8+ T cell responses have been conducted on the C57BL/6 (B6) background, limiting studies to MHC-K^b^ and D^b^ restricted responses. Therefore, our knowledge of CD8+ T cells restricted to other mouse MHC-1 molecules is limited.

The CD8+ T cell response to the intracellular parasite *Toxoplasma gondii (T. gondii)* differs dramatically between mouse strains, and determines whether or not mice eventually succumb to Toxoplasmic encephalitis. This difference is linked to MHC haplotype, with BALB/c (H-2^d^) mice being resistant and B6 (H-2^b^) mice being susceptible to infection. Resistance to *T. gondii* infection in H-2^d^ mice is due to an unusually strong and protective CD8+ T cell response directed against a 10-mer peptide (HF10) derived from the parasite antigen GRA6, presented by the MHC-1 molecule L^d^ (Brown et al. 1995; Blanchard et al. 2008). The GRA6-L^d^ specific CD8+ T cell response is maintained by a proliferative intermediate population that gives rise to large numbers of armed effector T cells throughout chronic infection, and shows no signs of functional exhaustion (Chu et al. 2016). In contrast, high levels of CD8+ T cell dysfunction have been reported in B6 mice after infection with *T. gondii* (Bhadra et al. 2011; Bhadra, Gigley, and Khan 2012). Interestingly, the MHC-1 L^d^ molecule shares key polymorphic residues with protective HLA-B alleles in HIV “elite controllers” which correlate with restricted peptide binding (Pereyra et al. 2010; Košmrlj et al. 2010; Narayanan and Kranz 2013).

Here we investigated the link between the atypical MHC-1 molecule L^d^ and the generation of “elite controller” CD8+ T cell responses. First, we compared the GRA6-L^d^ specific CD8+ cell response to the OVA-K^b^ specific response restricted to the conventional MHC-1 K^b^ molecule in their ability to protect against *T. gondii* infection. We also investigated the effect of a point mutation in MHC-1 L^d^ which leads to broader peptide binding, on another robust L^d^-restricted response during chronic Murine Cytomegalovirus (MCMV) infection (Pahl-Seibert et al. 2005). In both systems, T cells restricted to MHC-1 L^d^ exhibited resistance to exhaustion compared to T cells restricted to an MHC-1 with broader peptide binding. Overall, our results indicate that limited peptide binding by MHC correlates with the development of CD8+ T cell responses that resist exhaustion, and are consistent with a model in which limited peptide binding by MHC-1 and the corresponding low self-reactivity of CD8+ T cells favor protective responses during chronic infections.

## Results

### The CD8+ T cell response to GRA6-L^d^ is protective and resistant to exhaustion

We previously showed that the GRA6-L^d^ specific CD8+ T cell response to *T. gondii* infection is immunodominant and highly protective (Blanchard et al. 2008; Chu et al. 2016). To further evaluate the protective capacity of this T cell response, we performed a direct comparison with the well-characterized and potent CD8+ T cell response to OVA-K^b^ (Pope et al. 2001; Kumar and Tarleton 2001; Radford et al. 2002; Braaten et al. 2005; Miyakoda et al. 2012). We used TCR transgenic mice expressing rearranged transgenes representative of either the GRA6-L^d^ (TG6) or OVA-K^b^ (OT-1) response (Chu et al. 2016; Hogquist et al. 1994). Both TCRs exhibit similarly strong binding to their respective peptide-MHC complexes (Kd of 0.5-6 micromolar) ((Alam et al. 1999; Rosette et al. 2001) and data not shown). We also used a *T. gondii* strain in which both the model antigen OVA and the endogenous parasite antigen GRA6 are secreted from the parasite via dense granules (Schaeffer et al. 2009).

We first compared the protective capacity of TG6 and OT-1 T cells during *T. gondii* infection. We transferred naïve splenic TG6 or OT-1 CD8+ T cells from TCR transgenic H-2^b/d^ mice into T and B cell deficient H-2^b/d^ (B6 Rag2^-/-^ x BALB/c Rag2^-/-^) mice one day prior to infection, and monitored parasite load and T cell responses 6-8 weeks later **(Figure 1A)**. Mice that received OT-1 T cells had approximately 30-fold higher parasite load in the brain compared to mice that received TG6 T cells **(Figure 1B)**. T cell numbers in the brains of OT-1 transferred mice were higher than in TG6 transferred mice, implying that reduced protection was not due to reduced T cell expansion or trafficking to the brain **(Figure 1C)**. Interestingly, GRA6-L^d^ specific T cells could be readily detected by tetramer staining amongst donor T cells from OT-1 TCR transgenic mice (data not shown), suggesting that the outgrowth of GRA6-L^d^ specific T cells using endogenous TCR gene rearrangements could account for some protection in these mice. Consistent with this interpretation, we observed even more striking differences in cyst load, and a change in mouse survival, when splenocytes from TG6 Rag2-/- or OT-1 Rag2-/- mice were used as the donor populations **(Supplementary Figure 1)**.

**Figure 1:**
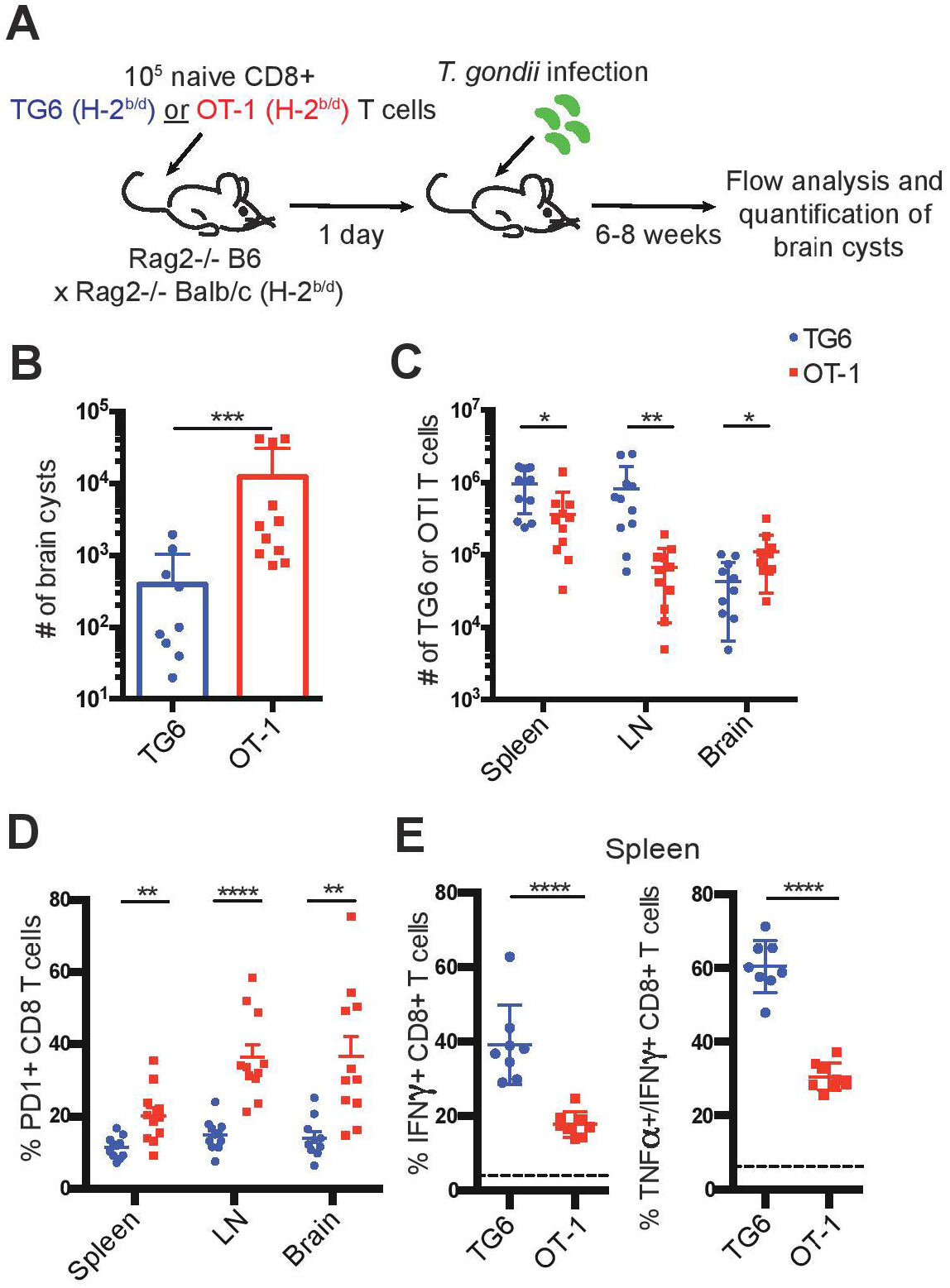
TG6 T cells are more protective and resistant to exhaustion compared to OT-1 T cells. **(A)** Experimental design to compare the protective capacity of TG6 or OT-1 T cells. B6 Rag2-/- x BALB/c Rag2-/- mice were injected intravenously (i.v.) with 10^5^ naïve CD8+ T cells from TG6 (n=11) or OT-1 (n=11) TCR transgenic mice, infected one day later with 2,000 *T. gondii* Pru-OVA parasites, and analyzed at 6-8 weeks post infection. **(B)** Number of brain cysts in mice transferred with TG6 (avg. 397 cysts) or OT-1 (avg. 12,453 cysts) CD8+ T cells. **(C)** Numbers of TG6 or OT-1 T cells recovered in the spleen, lymph nodes and brain determined by flow cytometry. **(D)** Frequency of PD1 expression on TG6 or OT-1 CD8+ T cells. **(E)** Production of IFNγ and TNFα cytokines by TG6 or OT-1 CD8+ T cells isolated from the spleen (n=8). Splenocytes were stimulated *ex vivo* with the HF10 (GRA6) or SIINFEKL (OVA) peptides in culture for 3 hours, and cells were analyzed for cytokine production by flow cytometry. Values indicate the percentage of IFNγ+ CD8+ T cells (left) or the percentage of TNFα producing cells among the IFNγ+ CD8+ T cells (right), following stimulation with peptide. Dotted line indicates background cytokine staining in un-stimulated cells from the same mice. Graphs show the summary of two independent experiments. Statistical significance was measured by a Mann-Whitney test in (B) and a t-test in (C)-(E) (*p<0.05, **p<0.01, ***p<0.001, and ****p<0.0001).

We also observed that OT-1 T cells recovered from chronically infected mice were functionally impaired compared to TG6 T cells. OT-1 T cells in the spleen, lymph nodes and brain displayed significantly higher expression of the T cell exhaustion marker PD1 **(Figure 1D)**. Additionally, OT-1 T cells in the spleen had a dramatically reduced capacity to produce the inflammatory cytokines IFNγ and TNFα compared to TG6 T cells **(Figure 1E)**.

The increase in T cell exhaustion in mice transferred with OT-1 versus TG6 T cells could be due differences in parasite load, and/or T cell intrinsic differences. To control for the effects of parasite load, we co-transferred both T cell populations into wild type H-2^b/d^ mice before infection with *T. gondii*. Again, we observed an increase in PD1 expression on OT-1 T cells compared to TG6 T cells in the spleen and lymph nodes of infected recipient mice **(Supplementary Figure 2)**. This difference was less dramatic than that seen in Rag2-/- mice, likely due to the effective parasite control by the GRA6-L^d^ specific T cells. These data suggest that T cell-intrinsic differences contribute to the difference in exhaustion between OT-1 and TG6 T cells.

To extend these observations to a polyclonal T cell response, we used fluorochrome-labeled GRA6-L^d^ and OVA-K^b^ tetramers to detect endogenous T cell responses in wild type H-2^b/d^ mice infected with OVA-expressing *T. gondii*. Both T cell populations expanded robustly in the spleen and lymph nodes of infected mice, although the GRA6-L^d^ specific population exhibited slightly better expansion, especially in the spleen **(Figure 2A-C)**. As expected, given the effective control of parasites in mice harboring the H-2^d^ allele, inhibitory receptors remained generally low on CD8+ T cells in these mice. However, in approximately half of the mice, OVA-K^b^ specific T cells expressed significantly elevated levels of the inhibitory receptors PD1, Tim3 and Lag3, compared to GRA6-L^d^ specific T cells in both the spleen and lymph nodes **(Figure 2D)**.

**Figure 2:**
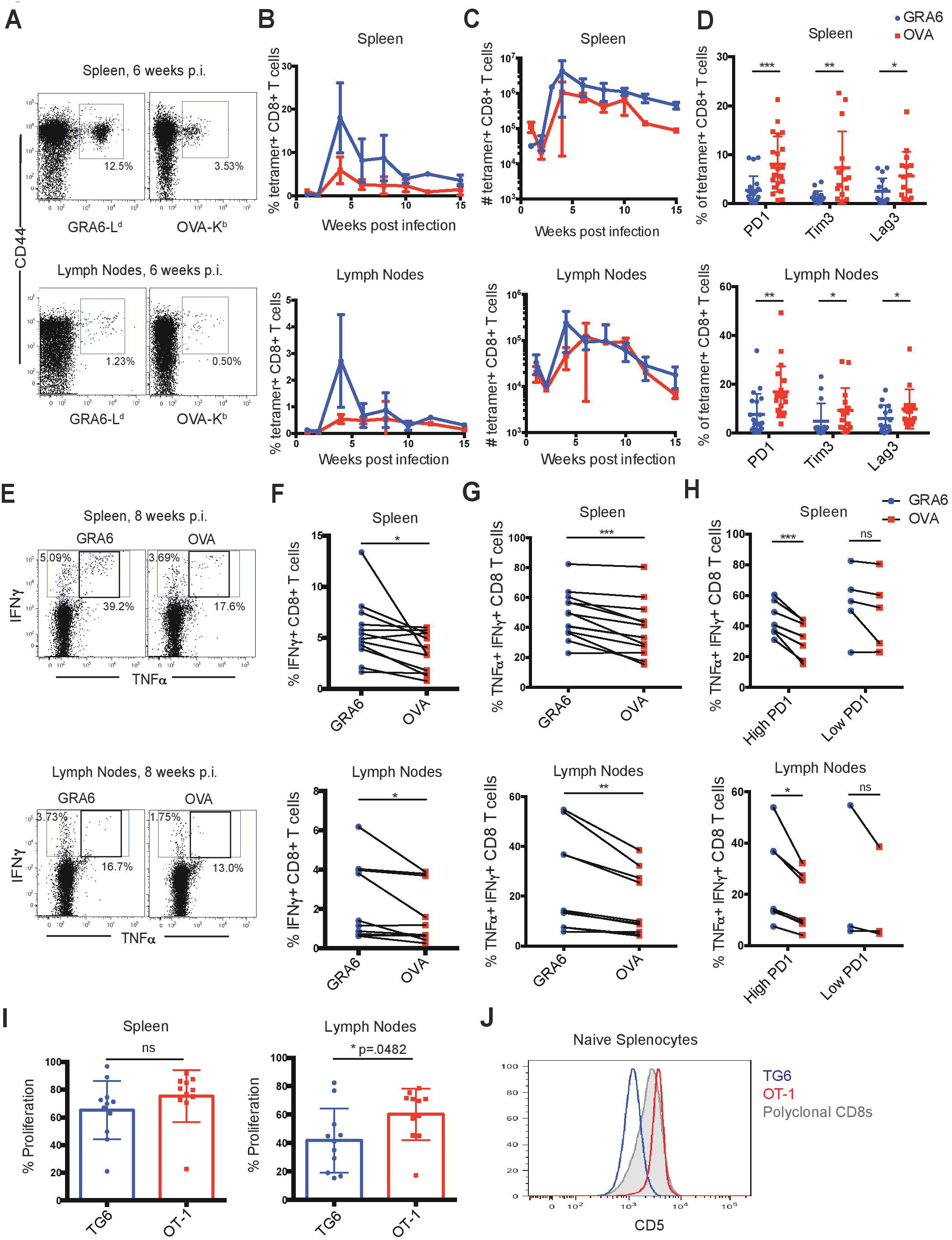
The OVA-K^b^ specific CD8+ T cell response to *T. gondii* infection shows signs of functional exhaustion. **(A-H)** H-2^b/d^ mice were infected intra-peritoneally (i.p.) with 10^5^ *T. gondii* Pru-OVA parasites and were sacrificed during the chronic infection (3-15 weeks). **(A)** Representative flow cytometry plots gated on live, singlet, CD8+ T cells and **(B)** compiled data showing percentage and **(C)** absolute numbers of GRA6-L^d^ (blue) and OVA-K^b^ (red) tetramer+ CD8+ T cells from the spleens and lymph nodes of infected mice. Graphs in (B)-(C) represent a total of 2-6 mice at each time point (n=26 (spleen), n=22 (lymph nodes)), and graphs display mean values with error bars depicting the SEM. **(D)** Expression of T cell exhaustion markers PD1, Tim3 and Lag3 on GRA6-L^d^ and OVA-K^b^ tetramer+ CD8+ T cells in the spleen and lymph nodes (n=21 (PD1), n=15 (Tim3/Lag3). **(E-H)** Production of IFNγ and TNFα cytokines by GRA6 or OVA specific T cells isolated from the spleens (n=13) or lymph nodes (n=10) of infected mice. Splenocytes or lymph node cells were stimulated *ex vivo* with the HF10 (GRA6) or SIINFEKL (OVA) peptides at 1μM in culture for 3 hours, and cells were analyzed for cytokine production by flow cytometry. **(E)** Representative flow cytometry plots of IFNγ and TNFα in CD8+ T cells from the spleen or lymph node. Compiled data showing the percentage of IFNγ+ cells among CD8+ T cells **(F)**, or the percentage of TNFα producing cells among the IFNγ+ CD8+ T cells in the spleen and lymph nodes **(G)**. **(H)** Data shown in (G) are separated based on PD1 expression in each mouse, with “high PD1” defined as >5% PD1+ in GRA6-L^d^ or OVA-K^b^ CD8+ T cells. Data points from the same animal are indicated by connecting lines. Graphs in (D) and (F-H) are the summary of >5 independent experiments, with data from all chronic time points (3-15 weeks) graphed together. **(I)** GRA6 and OVA antigen presentation in the spleens and lymph nodes of mice infected with *T. gondii* Pru-OVA. H-2^b/d^ mice were infected i.p. with 10^5^ *T. gondii* Pru-OVA parasites. At 3-8 weeks post infection, mice were injected i.v. with naïve CFSE-labeled splenocytes consisting of 10^6^ TG6 and 10^6^ OT-1 CD8+ T cells and analyzed by flow cytometry 3 days post T cell transfer. Transferred T cells were distinguished by a congenic marker together with Vβ2 (TG6) or Vβ5 (OT-1). Proliferation of TG6 and OT-1 CD8+ T cells was assessed by dilution of CFSE in the spleen and lymph nodes. Values are the percent proliferated cells out of the transferred T cell population. Graphs in (I) show compiled data from 4 independent experiments (n=11). **(J)** CD5 expression on splenic CD8+ T cells from TG6 (blue) and OT-1 (red) TCR transgenic mice. CD5 expression on polyclonal CD8+ T cells is shown for comparison (grey). Histogram is representative of >5 independent experiments. Statistical significance was measured by a t-test in (D, I) and a paired t-test in (F-H) with data from the same mouse paired (*p<0.05, **p<0.01, and ***p<0.001, ns is not significant).

To confirm that the expression of inhibitory receptors on OVA-K^b^ specific T cells correlates with reduced functionality of these cells, we analyzed cytokine production by GRA6-L^d^ and OVA-K^b^ specific T cells. We re-stimulated splenocytes or lymph node cells from infected mice *ex vivo* with GRA6 or OVA peptide and measured IFNγ and TNFα production by flow cytometry. Stimulation with the GRA6 peptide resulted in slightly higher proportion of IFNγ producing cells compared to the OVA peptide **(Figure 2E, F)**, which is likely a reflection of the relative size of the antigen-specific population **(Figure 2B)**. To assess the functionality of antigen-specific T cells on a per cell basis, we determined the percentage of IFNγ+ cells that also produced TNFα. This proportion was lower amongst OVA-K^b^ specific T cells, particularly in those mice with relatively high (>5%) PD1 expression on T cells **(Figure 2G, H)**. Thus, high PD1 expression correlated with reduced cytokine production in OVA-K^b^ specific T cells, indicating that OVA-K^b^ specific T cells are less functional compared to GRA6-L^d^ specific T cells from the same infected mice.

T cell exhaustion can be caused by chronic exposure to high levels of inflammatory signals or antigen (Wherry and Kurachi 2015). Differences in inflammatory signals are unlikely to account for the differences observed between GRA6-L^d^ and OVA-K^b^ responses when compared in the same infected mice. Thus we tested whether differences in antigen levels could account for these differences. While both GRA6 and OVA precursor proteins are secreted by dense granules during infection, differences in the expression levels or antigen processing efficiency could lead to differences in the level of peptide presentation (Gubbels et al. 2005; Gregg et al. 2011; Lopez et al. 2015; Buaillon et al. 2017; Tsitsiklis, Bangs, and Robey 2019). To test this possibility, we used an *in vivo* antigen presentation assay. We transferred equal numbers of naïve CFSE-labeled TG6 and OT-1 CD8+ T cells from TCR transgenic mice into chronically infected mice and assessed their proliferation by CFSE dilution 3 days after transfer. We observed a similar proportion of proliferating cells in TG6 and OT-1 T cells recovered from the spleen and lymph nodes of infected mice **(Figure 2I)**. As an additional measure of antigen stimulation, we also examined expression of Nur77, a TCR target gene whose induction is proportional to the strength of TCR stimulus (Moran et al. 2011). We did not observe a significant difference in Nur77 expression between TG6 and OT-1 T cells 3 days after transfer, suggesting TG6 and OT-1 T cells are receiving similar amounts of antigen stimulation **(Supplementary Figure 3)**. The observation that OVA-K^b^ specific T cells show more exhaustion than GRA6-L^d^ specific T cells, in spite of exposure to similar levels of antigen and inflammatory signals and a similar affinity for peptide-MHC, suggested that there may be intrinsic differences between these two T cell populations that influence their susceptibility to functional exhaustion.

### Limited peptide binding by L^d^ correlates with the selection of T cells with low self-reactivity

We considered that the unusual peptide binding characteristics of MHC-1 L^d^ might play a role in generating exhaustion-resistant T cell responses. Expression of L^d^ on the cell surface is 2- to 3-fold lower than that of other conventional MHC molecules, and several groups have reported that this is due to poor binding to endogenous (self) peptide ligands (Potter et al. 1981; Lie et al. 1991; Narayanan and Kranz 2013). Since L^d^ binds poorly to self-peptides, we might expect that positive selection on L^d^ would produce T cells with relatively low self-reactivity. To address this possibility, we examined expression of CD5, a robust surrogate marker for self-reactivity (Azzam et al. 2001; Fulton et al. 2015; Persaud et al. 2014) on T cells from TG6 transgenic mice. We found that CD5 surface expression on naïve splenocytes from TG6 TCR transgenic mice is relatively low compared to polyclonal CD8+ T cells **(Figure 2J)**, supporting that TG6 T cells are less self-reactive than polyclonal CD8+ T cells. On the other hand, splenocytes from OT-1 TCR transgenic mice, which are selected on the conventional MHC molecule K^b^, have relatively high CD5 expression, consistent with previously published literature that OT-1 T cells have high self-reactivity (Ge et al. 2004) **(Figure 2J)**.

### The L^d^-restricted MCMV IE1 specific CD8+ T cell response has a proliferative CXCR3+KLRG1+ intermediate population

Given the unusual properties of L^d^, we considered whether other L^d^ restricted CD8+ T cell responses might also be particularly robust during chronic infection. The IE1-L^d^ specific CD8+ T cell response to MCMV is one of a handful of “inflationary” responses that expands during chronic infection and provides strong protection (Pahl-Seibert et al. 2005; Rafaela Holtappels et al. 2002). During *T. gondii* infection, the GRA6-L^d^ CD8+ T cell response is maintained by an intermediate population (Tint), defined by dual expression of CXCR3 and KLRG1, which is highly proliferative and replenishes a short-lived effector (Teff) population to allow for a long-lasting effector T cell response (Chu et al. 2016). Interestingly, we observed the same Tint population amongst IE1-L^d^ specific T cells in mice chronically infected with MCMV, whereas this population was less prominent in CD44+ tetramer-CD8+ T cells from the same mice **(Figure 3A, B)**. The Tint population was also the most proliferative population within IE1-L^d^ specific T cells, as measured by Ki67 expression **(Figure 3C)**, similar to what we previously observed in the GRA6-L^d^ specific response (Chu et al. 2016). Together, these data suggest that two robust L^d^-restricted responses may be similarly maintained during chronic infection.

**Figure 3:**
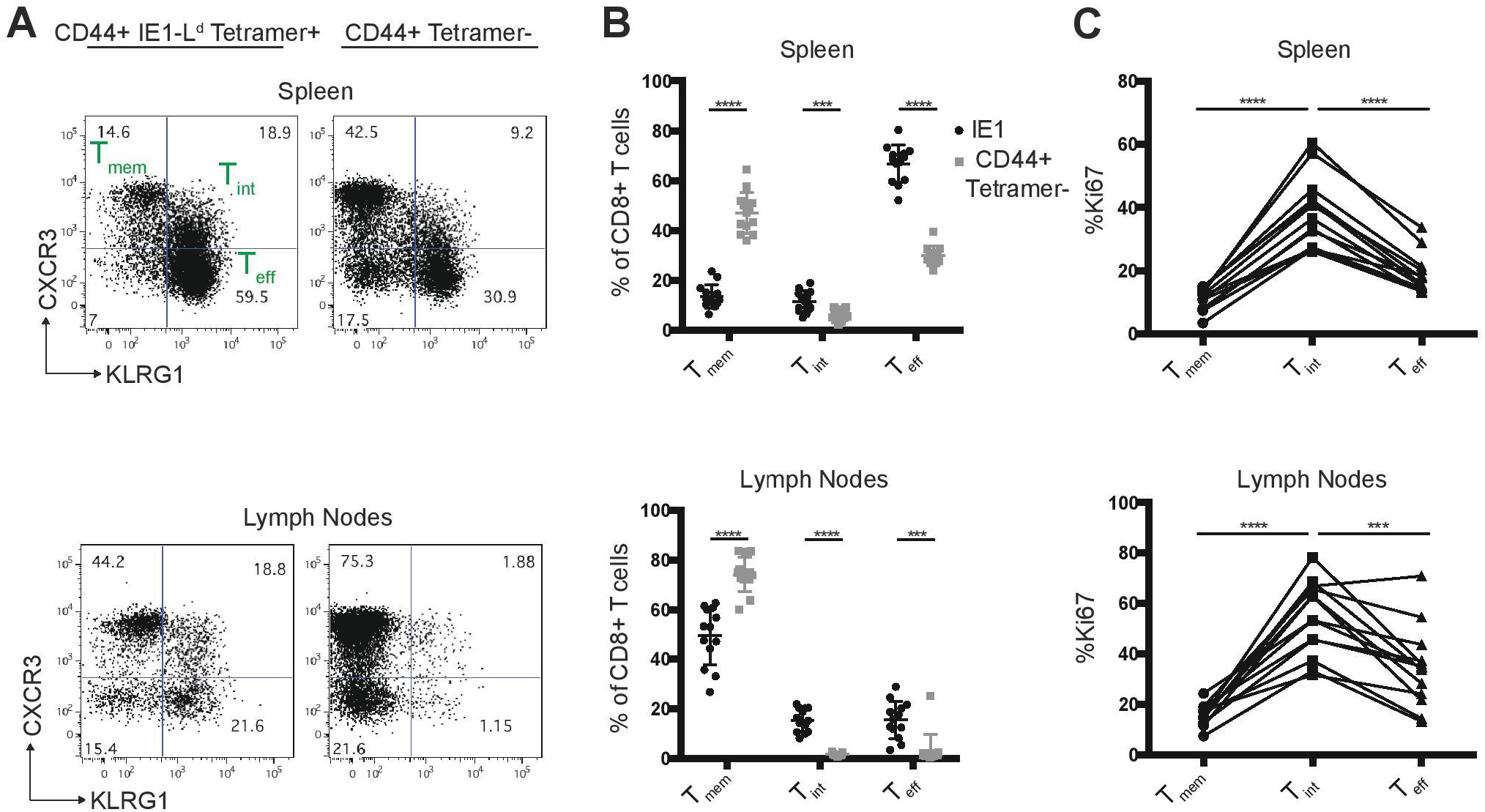
The L^d^-restricted MCMV IE1 response has a proliferative intermediate population. H-2^d/d^ mice were infected i.p. with 10^6^ pfu MCMV and cells from the spleens and lymph nodes were analyzed by flow cytometry 11-16 weeks post infection. **(A)** Representative flow cytometry plots show CXCR3 and KLRG-1 expression within CD44+ IE1-L^d^ and CD44+ tetramer-CD8+ T cells in the spleens and lymph nodes of MCMV infected mice. **(B)** Percentage of CXCR3+KLRG1- (Tmem), CXCR3+KLRG-1+ (Tint), and CXCR3-KLRG-1+ (Teff) cells within IE1-L^d^ (black) and CD44+ tetramer-(grey) CD8+ T cells. **(C)** Percentage of Ki67+ cells within the indicated subset defined by CXCR3 and KLRG-1 expression in IE1-L^d^ CD8+ cells in the spleens and lymph nodes. Data points from the same animal are indicated by connecting lines. Graphs are the summary of 3 independent experiments (n=13). Statistical significance was determined by a t-test in (B) and a oneway ANOVA in (C) (*p<0.05, **p<0.01, ***p<0.001, and ****p<0.0001).

### An H2-L^d^ variant with broader peptide binding is associated with increased expression of exhaustion markers by IE1-L^d^ specific T cells during MCMV infection

We next wanted to investigate whether limited or broad peptide binding in the context of H2-L could contribute to the exhausted T cell phenotype. Previous studies in cell lines have shown that an HIV-controller associated polymorphism at amino acid position 97 in the mouse H2-L gene impacts the broadness of peptide binding (R. Smith et al. 2002; Narayanan and Kranz 2013). To extend these studies to an *in vivo* setting, we used CRISPR-EZ (Chen et al. 2016) to generate mice in which the controller-associated tryptophan at position 97 of L^d^ is converted to a progressor-associated arginine residue (L^d^ W97R mutation) **(Figure 4A)** (Pereyra et al. 2010). As expected based on studies in cell lines (Narayanan and Kranz 2013), splenocytes from L^d^ 97R mice have higher surface expression compared to L^d^ 97W, consistent with the enhanced binding to self-peptides **(Figure 4B)**. Additionally, while incubation with an antigenic peptide stabilized wild type (97W) L^d^, this had little impact on the levels of 97R L^d^ **(Figure 4D-E)**. In some cell types (e.g. CD4+ T cells) the levels of IE1 peptide-stabilized L^d^ 97W were equivalent to L^d^ 97R **(Supplementary Figure 4)**. Thus, limited self-peptide binding by L^d^ 97W is reversed by the L^d^ W97R mutation, allowing for higher cell surface expression.

**Figure 4:**
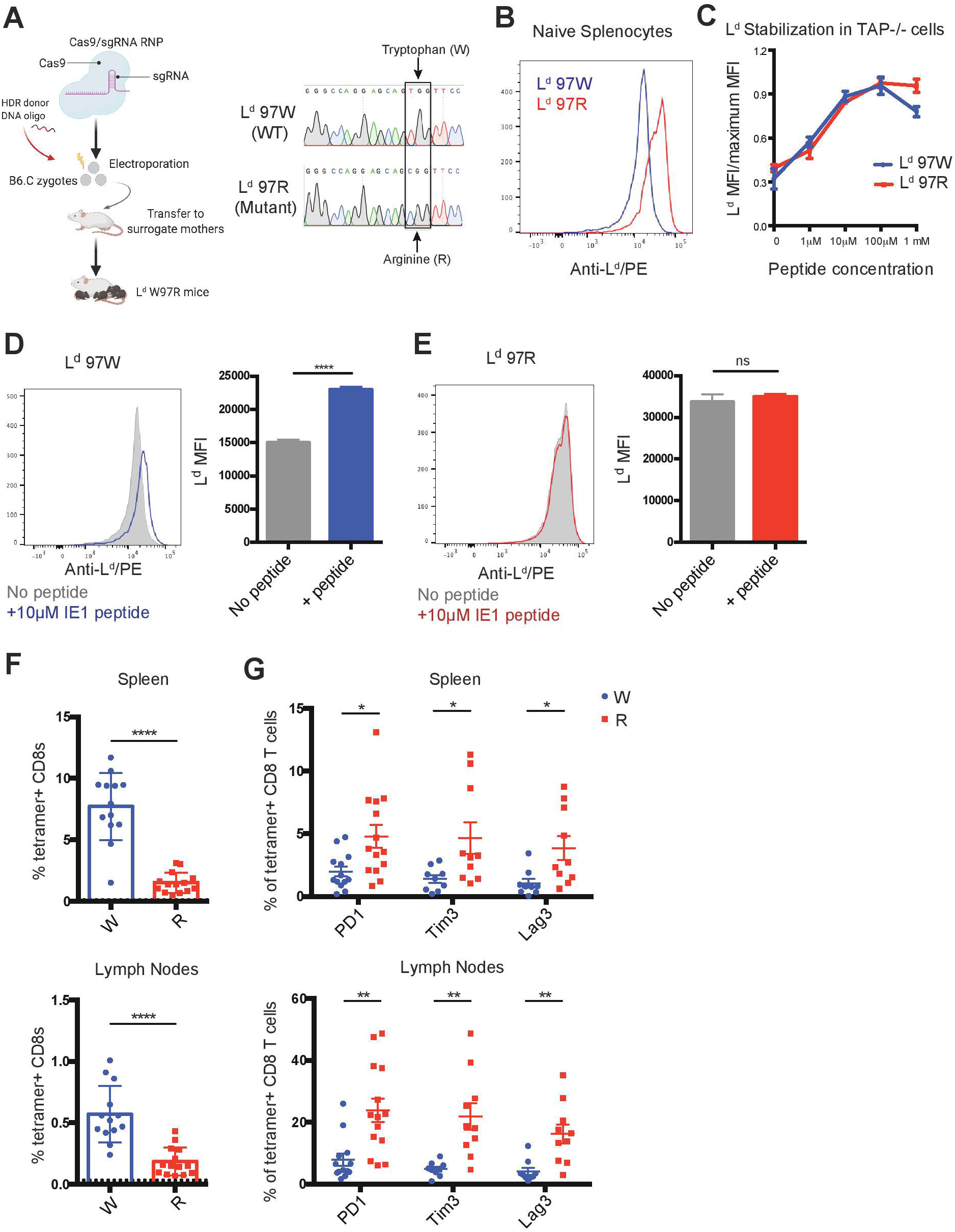
Polymorphism in H2-L^d^ increases self-peptide binding and correlates with increased expression of exhaustion markers on IE1-L^d^ specific CD8+ T cells. **(A)** Generation of L^d^ W97R mutant mouse by CRISPR-EZ (Chen et al. 2016; Modzelewski et al. 2018). Cas9-sgRNA ribonucleoprotein (RNP) was prepared and incorporated together with the HDR donor DNA oligo and electroporated into mouse zygotes from H-2^d^ (B6.C) mice. A single T→C point mutation that converts a tryptophan (TGG) to arginine (CGG) at position 97 in the L^d^ gene was generated. Sequences show an example of a wild-type B6.C mouse compared to an L^d^ W97R mutant mouse generated by sequencing DNA from the tail. **(B)** Surface expression of H2-L^d^ on naïve splenocytes from L^d^ W97 (blue) or L^d^ R97 mice (red). H2-L^d^ surface expression was measured by flow cytometry using an antibody that binds peptide-bound L^d^. Data is representative of 3 independent experiments. **(C)** IE1 peptide binding strength to H-2L^d^. Binding strength was measured using the ability of peptide to stabilize surface L^d^ expression on the TAP deficient RMA-S cell line expressing either the WT (97W) or mutated (97R) version of H2-L^d^. The mean fluorescent intensity (MFI) of peptide-bound H2-L^d^ was measured after incubating the cells with increasing concentrations of peptide for 3 hours. Graph shows representative data from 2 independent experiments. **(D-E)** Surface expression of H2-L^d^ on naïve splenocytes from L^d^ W97 (D) or L^d^ R97 (E) mice with (color) or without (grey) the addition of 10μM IE1 peptide. Experimental groups consisted of 3 samples, and data is representative of 2 independent experiments. **(F-G)** L^d^ W97 (n=13) or L^d^ R97 (n=14) mice were infected with 10^6^ pfu MCMV and sacrificed at 11-16 weeks post infection. **(F)** IE1-L^d^ tetramer+ CD8+ T cells were quantified in the spleen and lymph nodes of infected mice. Dotted line indicates background IE1-L^d^ tetramer staining in CD4+ T cells. **(G)** Expression of exhaustion markers PD1, Tim3, and Lag3 on IE1-L^d^ tetramer+ CD8+ T cells in the spleen and lymph nodes in L^d^ W97 and R97 mice. Graphs are pooled data from 3 independent experiments. Statistical significance was determined by a t-test (*p<0.05, **p<0.01, and ****p<0.0001).

To further explore the connection between limited peptide binding by MHC and robust CD8+ T cell responses, we investigated the effect of the L^d^ W97R mutation in the setting of persistent infection. We focused on the MCMV IE1-L^d^ response, since this peptide, unlike the *T. gondii* GRA6 peptide, binds well to both 97R and 97W versions of L^d^ (**Figure 4C** and data not shown) (Narayanan and Kranz 2013). We infected L^d^ 97W and L^d^ 97R mice with MCMV and characterized the IE1-L^d^ specific CD8+ T cell response during chronic infection (11-16 weeks). IE1-L^d^ specific T cells expand robustly in L^d^ 97W mice, while expansion is weaker but still readily detectable in L^d^ 97R mice **(Figure 4F)**. Furthermore, expression of the exhaustion markers PD1, Tim3 and Lag3 is significantly higher on IE1-L^d^ specific T cells in L^d^ 97R mice, in both the spleen and lymph nodes **(Figure 4G)**. Interestingly, expression of exhaustion markers and the activation marker CD69 is markedly higher in the lymph nodes compared to the spleen in both L^d^ 97W and L^d^ 97R mice **(Figure 4G; Supplementary Figure 5)**, although the IE1-L^d^ specific T cell population is smaller in the lymph nodes. This may be a result of higher antigen load in the lymph nodes during MCMV infection (Karrer et al. 2003; Torti et al. 2011), possibly pushing IE1-L^d^ specific T cells in the lymph nodes towards exhaustion and eventual deletion.

MCMV infected L^d^ 97W and L^d^ 97R mice have similar viral loads in the salivary glands **(Supplementary Figure 6A)**, indicating that the difference in the IE1-L^d^ specific response is unlikely to be due to a difference in antigen load. To confirm that the changes in IE1-L^d^ specific T cells are a direct consequence of the W97R mutation, we examined another immunodominant response restricted to a different MHC-1 molecule (m164-D^d^) (R. Holtappels et al. 2002). In contrast to IE1-L^d^ specific T cells, m164-D^d^ specific T cells do not exhibit an increase in expression of PD1, CD69, and Nur77 **(Supplementary Figure 6B-D)**. Therefore, these data support our hypothesis that an intrinsic difference between T cells selected on MHCs with limited versus broad peptide binding may contribute to resistance to exhaustion.

## Discussion

While MHC-1 molecules with limited peptide binding are associated with HIV control in humans and *T. gondii* control in mice, the link between the peptide binding characteristics of MHC and effective CD8+ T cell responses has not been experimentally examined. We used two parallel approaches to address this issue. First, we compared two high affinity CD8+ T cell responses, one restricted to an MHC molecule with limited peptide binding (L^d^), and another restricted to a typical MHC-1 molecule (K^b^) in the same *T. gondii*-infected mice. Second, we compared T cell responses to the same viral peptide presented by either L^d^, or a single amino acid variant of L^d^ with broad peptide binding, during chronic MCMV infection. In both cases, T cells restricted to MHC-1 L^d^ showed fewer signs of exhaustion compared to T cells restricted to MHC-1 with broader binding to self-peptides. These data suggest that MHC molecules with limited peptide binding allow for the development of T cells with the ability to persist during chronic infection without becoming exhausted.

At first glance, the association between limited peptide binding and highly protective CD8+ T cell responses appears contradictory. However, while elite control associated MHC molecules bind less broadly to peptides in general, they nevertheless bind strongly to certain antigenic peptides (Narayanan and Kranz 2013; Schneidewind et al. 2008; Košmrlj et al. 2010). Importantly, MHC-1 L^d^ binds poorly to self-peptides, leading to lower cell surface expression and reduced stability of surface peptide-MHC complexes (Potter et al. 1981; Lie et al. 1991; Narayanan and Kranz 2013). The inability of L^d^ to bind strongly to self-peptides may explain the unusually low self-reactivity of TG6, a public TCR representative of the CD8+ T cell response to the dominant *T. gondii* antigen GRA6 (data not shown). These data suggest that the combination of low reactivity to self, coupled with high reactivity to a pathogen-derived peptide may be a key feature of elite controller CD8+ T cell responses.

These considerations support a model in which MHC-1 molecules with restricted peptide binding properties may be considered “specialists”, in contrast to the majority of “generalist” MHC-1 molecules with broad peptide binding **(Supplementary Figure 7)** (Chappell et al. 2015). During thymic development, generalist MHC molecules select T cells with relatively high self-reactivity, whereas specialist MHC molecules select T cells with relatively low self-reactivity due to weak binding to self-peptides **(Supplementary Figure 7A, B).** While most infections favor expansion of a broad set of T cells with relatively high self-reactivity, in settings where pathogen-derived peptides are able to bind to, and stabilize, specialist MHC-1 molecules, T cells with high reactivity to a pathogen-derived peptide, but low reactivity to self can expand, giving rise to elite controller CD8+ T cell responses **(Supplementary Figure 7C, D).**

Why might low self-reactivity favor elite controller T cell responses? As suggested previously (Košmrlj et al. 2010), T cells selected in the thymus on specialist MHC molecules may experience less stringent negative selection, allowing for the development of unusually high affinity T cells, which would have been deleted by a generalist MHC, to enter the mature T cell repertoire. Specifically, while negative selection primarily functions to remove highly cross-reactive thymocytes (Huseby et al. 2005, 2006; McDonald et al. 2015), modeling studies have predicted that limited self-peptide presentation by MHC allows for the selection of T cells with high cross-reactivity (Košmrlj et al. 2010). In support of this prediction, an MHC-2 molecule presenting a single peptide selects T cells with highly cross-reactive TCRs by taking advantage of high affinity germline interactions, as a result of minimal negative selection (Dai et al. 2008).

In addition to the selection of a distinctive TCR repertoire, specialist MHC molecules may also impact the functional tuning of T cells during their development in the thymus. Recent studies have shown that T cells tune their functional responsiveness based on their degree of self-reactivity, with higher self-reactivity (CD5^high^) T cells exhibiting greater IL-2 production, more rapid expansion during acute infection, and greater responsiveness to cytokines (Weber et al. 2012; Mandl et al. 2013; Persaud et al. 2014; Fulton et al. 2015; Cho et al. 2016; Hogquist and Jameson 2014). While CD5^high^ T cells are generally superior in their responses to pathogens, in one study CD5^low^ CD4+ T cells were relatively resistant to activation induced cell death, and responded better upon secondary challenge (Weber et al. 2012). It is tempting to speculate that the reduced responsiveness of CD5^low^ T cells to inflammatory environments may allow them to better avoid exhaustion during chronic infection.

In addition to functional tuning during T cell selection in the thymus, the degree of self-reactivity may also impact the tonic TCR signals that T cells experience in the periphery. Tonic signals have been shown to contribute to inhibitory receptor upregulation and functional hyporesponsiveness in the absence of antigen-specific stimulation (Hsu et al. 2017; Ellestad et al. 2017). Recently, Duong et al. demonstrated that TCRs with higher affinity and higher CD5 expression were more susceptible to functional hyporesponsiveness after chronic exposure to self-peptide MHC (Duong et al. 2019). Higher tonic signaling may push CD5^high^ T cells towards exhaustion, while CD5^low^ T cells restricted to the specialist MHC-1 molecule L^d^ may experience lower levels of tonic signaling, contributing to their resistance to exhaustion during chronic infection.

While our model predicts that T cells restricted to specialist MHC molecules have the potential to generate high affinity responses to pathogen-derived peptides, it is unlikely that TCR signal strength alone can account for the lack of functional exhaustion in the L^d^ restricted responses examined in this study. TG6 and OT-1 TCRs have similarly high affinity for antigen-MHC ((Alam et al. 1999; Rosette et al. 2001) and data not shown) and show similar levels of the TCR signal strength reporter Nur77 in infected mice. Likewise IE1-L^d^ specific T cells in L^d^ 97R and L^d^ 97W mice show comparable functional avidity based on peptide titration experiments (data not shown). Moreover, PD1 induction and functional exhaustion is associated with strong, persistent TCR signals, implying that stronger TCR reactivity would lead to more, not less, exhaustion. Thus it seems likely that differences in functional tuning and tonic signals contribute, at least in part, to the propensity of T cells to become exhausted.

In addition to the T cell intrinsic factors explored here, it is clear that extrinsic factors such as persistent inflammatory signals and high antigen levels also play a role in driving T cell exhaustion. In the *T. gondii*-mouse infection model, the highly protective GRA6-L^d^ specific T cell response may contribute to a feedback mechanism to prevent exhaustion of other T cell responses by keeping antigen and inflammation levels low. Thus the OVA-K^b^ specific response exhibits less exhaustion in H-2^b/d^ compared to H-2^b^ mice due the ongoing GRA6-L^d^ specific response and reduced parasite load. On the other hand, MCMV is well controlled in mice, even in the absence of a particularly protective T cell response, which may explain the overall low level of T cell exhaustion in this model. Finally, for infections where pathogen levels and inflammation remain high in spite of strong T cell responses, extrinsic factors may be able to overcome the intrinsic resistance of certain T cells to exhaustion. Consistent with this notion, the L^d^-restricted CD8+ T cell response to NP peptide during mouse infection with LCMV clone 13 infection exhibited high levels of PD1 and functional exhaustion (data not shown).

In summary, we have used two experimental systems to demonstrate that limited peptide binding by MHC-1 L^d^ is associated with the generation of exhaustion-resistant CD8+ T cell responses. T cell intrinsic factors linked to the peptide binding properties of restricting MHC molecules may combine with T cell extrinsic factors, such as high antigen levels and persistent inflammatory signals, to determine the outcome of infection. Future studies should directly explore the role of TCR repertoire, T cell tuning, and tonic TCR signaling in T cell susceptibility to exhaustion. Furthermore, extensions to human studies may help to identify additional MHC-1 specialist molecules, and may provide insight into protective responses to HIV and other persistent infections, which would have valuable therapeutic implications.

## Materials and Methods

### Mice

B6 (C57BL/6), B6 Rag2-/- (B6-Cg-Rag2^*tm1.1Cgn*^/J), B6.C (B6.C-*H2d*/bByJ), BALB/c, BALB/c Rag2-/- (C.B6(Cg)-Rag2 ^*tm1.1Cgn*^/J), and TCR transgenic mice specific for OVA- K^b^ (OT-1; C57BL/6-Tg(TcraTcrb)1100Mjb/J) were originally purchased from The Jackson Laboratory (Bar Harbor, ME, USA) and then bred in the AAALAC accredited animal facility at University of California, Berkeley. TCR transgenic mice specific for GRA6-L^d^ (TG6) were previously generated in our lab as described (Chu et al. 2016). F1 mice (B6xB6.C or B6 Rag2-/- x BALB/c Rag2-/-) were used for all *T. gondii* experiments in order to monitor multiple *T. gondii* epitopes. L^d^ W97R mutant mice were generated in our lab (described below). Mice were infected at 6-8 weeks old, and sacrificed at indicated times post infection. All mouse procedures were approved by the Animal Care and Use Committee (ACUC) of the University of California.

### Infection

For *T. gondii* infections, C57BL/6 Rag2-/- x BALB/c Rag2-/- (H-2^b/d^) or intact C57BL/6 x B6.C (H-2^b/d^) mice were injected intraperitoneally (i.p.) with 2,000 or 10^5^ live tachyzoites of the type II Prugniuad-tomato-OVA strain (Pru-OVA) *T. gondii* (Schaeffer et al. 2009), respectively. *T. gondii* brain cysts were quantified by imaging of homogenized brain samples stained with fluorescein-conjugated *Dolichos biflorus* agglutinin (Vector Laboratories) to stain the cell wall with a protocol adapted from (S.-K. Kim and Boothroyd 2005). For MCMV infections, L^d^ 97W or 97R B6.C (H-2^d^) mice were injected i.p. with 10^6^ pfu Smith strain MCMV (L. M. Smith et al. 2008). MCMV viral titers were measured using quantitative PCR.

### Flow Cytometry

Cells were stained with various fluorophore-labeled antibodies from BD Biosciences (San Diego, CA), Biolegend (San Diego, CA), or Tonbo Biosciences (San Diego, CA) at 4°C. For intracellular staining, cells were fixed and permeabilized using the eBioscience FoxP3 Fixation/permeabilization buffer kit (for Nur77 and Ki67) or the BD Biosciences Cytofix/Cytoperm kit (for cytokines) according to manufacturer protocols. Biotinylated peptide-MHC monomers were obtained from the NIH Tetramer Facility (Atlanta, GA). Tetramers were made by conjugating the biotin-labeled monomers with PE-labeled or APC-labeled streptavidin (Agilent) according to protocols from the NIH tetramer facility. A tetramer with the WR97 mutation in L^d^ was generated by the NIH Tetramer Facility and used for staining the IE1-L^d^ tetramer specific populations in 11/14 of the L^d^ 97R mice. No significant differences were observed when using the L^d^ 97W versus 97R tetramer. Data were acquired on BD LSR Fortessa and analyzed using FlowJo software (Ashland, OR). Statistical analyses were conducted using GraphPad Prism (San Diego, CA).

### *Ex Vivo* Stimulation with Peptides for Quantification of Cytokine Production

Spleen or lymph node cells from infected mice were plated at a density of 1-3×10^5^ cells per well in 96-well round-bottom tissue culture plates and incubated with 1μM of the indicated peptide (Peptide 2.0 Inc.) in media for 3 hours in the presence of protein transport inhibitor cocktail (eBioscience). Cells were then harvested for surface and intracellular antibody staining before flow cytometry analysis.

### Antigen Presentation Detection *in Vivo*

Lymph nodes and spleen of TG6 (Ly5.2/5.2) or OT-1 (Ly5.1/5.2) TCR transgenic (H-2^b/d^) mice were harvested and cells were labeled with the cell proliferation dye CFSE (5μM). Cells were washed twice with RPMI and twice with PBS. Cells were then counted and mixed in equal proportions prior to transfer. 5×10^5^ TG6 and 5×10^5^ OT-1 cells were transferred by intravenous tail-vein injection into previously infected mice (Ly5.1/5.1) and proliferation dye dilution was examined 3 days after transfer by gating on congenic markers to identify each donor population. Relative antigen presentation is expressed as the percentage of donor TG6 or OT-1 cells that diluted the proliferation dye.

### Generation of L^d^ W97R mutant mouse by CRISPR/Cas9

Single guide RNAs (sgRNAs) were designed using MIT’s CRISPR RNA guide design tool (crispr.mit.edu). A homology directed repair (HDR) donor DNA oligo with the desired mutation was synthesized by Life Technologies Co., with a single T→C point mutation that converts a tryptophan (TGG) to arginine (CGG) at position 97 in the L^d^ gene. Cas9-sgRNA RNP was prepared and incorporated together with the HDR donor DNA oligo and electroporated into mouse zygotes from H-2^d^ (B6.C) mice as described (Chen et al. 2016; Modzelewski et al. 2018). Resulting mice were genotyped by sequencing. Mice were backcrossed to WT B6.C mice for several generations.

### MHC-1 stabilization assay

RMA-S cells were infected with the VSV.G plasmid expressing the 97W or 97R version of L^d^ and sorted for L^d^-positive cells using the Thy1.1 marker. The MHC-1 stabilization assay was performed as described (Feliu et al. 2013). In brief, RMA-S.L^d^ (97W or 97R) cells or splenocytes from L^d^ 97W and 97R mice were incubated at room temperature overnight to increase surface MHC-1 expression. The next day, cells were washed with PBS and plated at 3×10^5^ cells per well in a 96-well round-bottom tissue culture plate. Peptides of interest (Peptide 2.0 Inc.) were added to the cells at indicated concentrations. The cells were incubated for 1 hour at room temperature and 3 hours at 37°C. Cells were stained with the 30-5-7 antibody (specific for conformed, peptide-bound L^d^) and a goat anti-mouse IgG phycoerythrin (PE)-conjugated secondary antibody and analyzed by flow cytometry.

## Conflict of Interest

The authors declare that the research was conducted in the absence of any commercial or financial relationships that could be construed as a potential conflict of interest.

## Author Contributions

AT designed the research, conducted experiments, analyzed the data, and wrote the manuscript. ER designed the research and wrote the manuscript. DB, SC, KG, LLB, and YW conducted experiments. AT, LL and AM contributed to generation of the L^d^ W97R mutant mouse. EZ and SD designed the research.

## Funding

Funding was provided by National Institutes of Health (RO1AI065537 and AI093132) to EAR, and a Cancer Research Coordinating Committee Fellowship (AT).

## Acknowledgements

We would like to thank Angus Lee and Harman Dhaliwal (UC Berkeley Cancer Research Lab, Gene Targeting Facility) for their help in generating the L^d^ W97R mutant mice.

