## Supplementary Figures for "An unusual MHC molecule generates protective CD8+ T cell responses to chronic infection"

### Supplementary Material

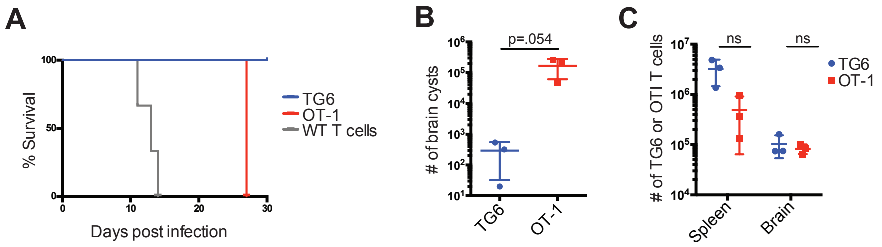


**Supplementary Figure 1.** Protective capacity of TG6 vs. OT-1 T cells *in vivo*. B6 Rag2-/- x BALB/C Rag2-/- (H-$2^{b/d}$) mice were injected i.v. with $10^6$ naïve TG6 Rag2-/- (n=6), OT-1 Rag2-/- (n=3) or wild type B6xBALB/c (n=3) splenocytes. The next day, mice were infected i.p. with 2,000 *T. gondii* Pru-OVA parasites, and then monitored for signs of illness and analyzed at day 27 post infection. **(A)** Survival curve indicating when mice were sacrificed due to illness. TG6 transferred mice remained healthy and were sacrificed together with OT-1 transferred mice for analysis. Survival of OT-1 transferred mice was significantly different from TG6 transferred mice (p=0.0047). **(B)** The number of brain cysts measured at 27 days post infection. **(C)** Numbers of TG6 or OT-1 CD8+ T cells recovered from the spleen and brain of recipient mice as measured by flow cytometry. Statistical significance was determined by a Gehan-Breslow-Wilcoxon test in (A), and a t-test in (B) and (C) (*p<0.05, **p<0.01, ns is not significant).

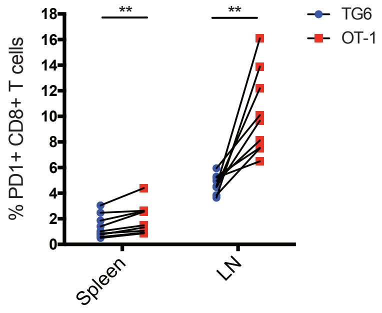


**Supplementary Figure 2.** H-$2^{b/d}$ mice were injected i.v. with splenocytes consisting of $10^6$ TG6 and $10^6$ OT-1 CD8+ T cells from TCR transgenic mice. Mice were infected i.p. with $10^5$ OVA-expressing *T. gondii* parasites the next day. Expression of PD1 on TG6 or OT-1 CD8+ T cells in the spleen and lymph nodes was quantified 6-12 weeks post infection (n=9). Data points from the same mouse are indicated by connecting lines. Statistical significance was determined by a paired t-test, with data from the same mouse paired (**p<0.01).

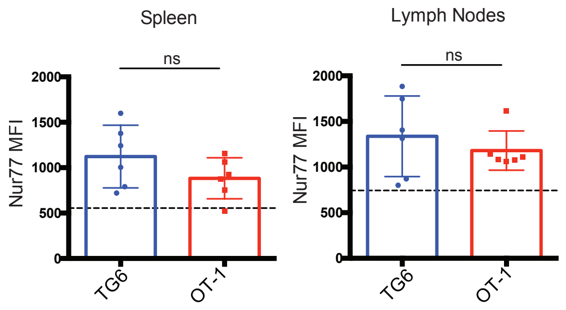


**Supplementary Figure 3.** H-$2^{b/d}$ mice were infected i.p. with $10^5$ *T. gondii* Pru-OVA parasites. At 3-8 weeks post infection, mice were injected i.v. with naïve CFSE-labeled splenocytes consisting of $10^6$ TG6 and $10^6$ OT-1 CD8+ T cells and analyzed by flow cytometry 3 days post T cell transfer. Transferred T cells were distinguished by a congenic marker, as well as either Vβ2 (for TG6) or Vβ5 (for OT-1). Graphs show mean fluorescence intensity (MFI) of intracellular Nur77 on CD44+ TG6 and CD44+ OT-1 CD8+ T cells in the spleen and lymph nodes. Dotted line indicates Nur77 expression in CD44- CD8+ cells from the same samples. Data are compiled from 2 independent experiments (n=6). Statistical significance was measured by a t-test (*p<0.05).

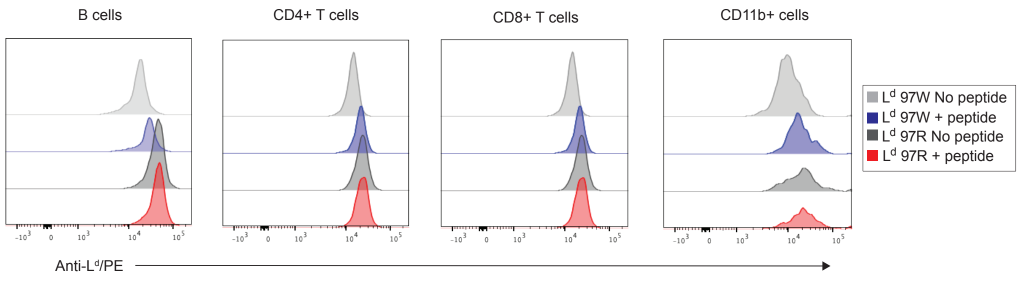


**Supplementary Figure 4.** Surface expression of H2-$L^d$ on **(A)** CD4+ T cells, **(B)** CD8+ T cells, **(C)** B cells, or **(D)** macrophages from splenocytes of $L^d$ W97 or $L^d$ R97 mice with or without the addition of 10μM IE1 peptide. Histograms are representative of 2 independent experiments.

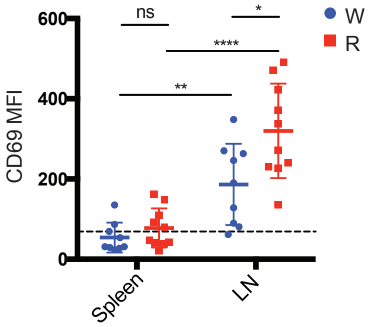


**Supplementary Figure 5.** L$^d$ 97W or L$^d$ 97R mice were infected with $10^6$ Pfu MCMV and sacrificed at 11-16 weeks post infection. CD69 expression on IE1-L$^d$ specific CD8+ T cells was quantified by flow cytometry. Dotted line indicates CD69 expression in CD44- CD8+ T cells in the same mice. Data is compiled from 2 independent experiments. Statistical significance was determined by a t-test (*p<0.05, **p<0.01, ***p<0.001, and ****p<0.0001, ns is not significant).

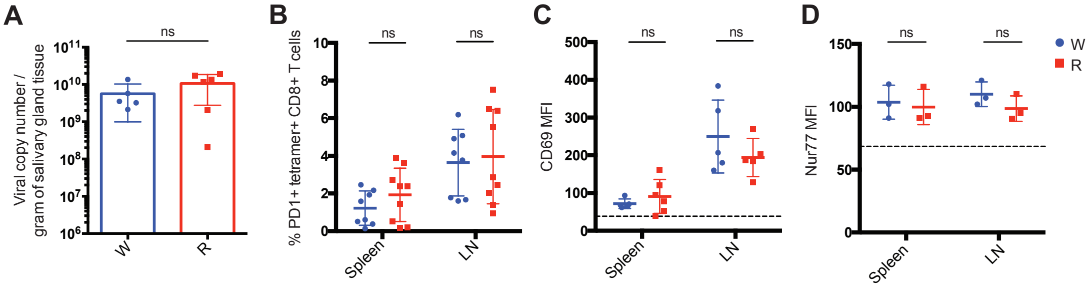


**Supplementary Figure 6.** L$^d$ 97W or L$^d$ 97R mice were infected with $10^6$ Pfu MCMV and sacrificed at 11-16 weeks post infection. **(A)** MCMV viral copy number in the salivary glands of L$^d$ 97W and L$^d$ 97R mice 14 weeks post infection was quantified by qPCR. Expression of **(B)** PD1, **(C)** CD69, and **(D)** Nur77 on m164-D$^d$ specific CD8+ T cells quantified by intracellular staining and flow cytometry. Dotted lines in (C-D) indicate CD69 or Nur77 expression in CD44- CD8+ T cells in the same mice. Statistical significance was determined by a t-test (*p<0.05, ns is not significant).

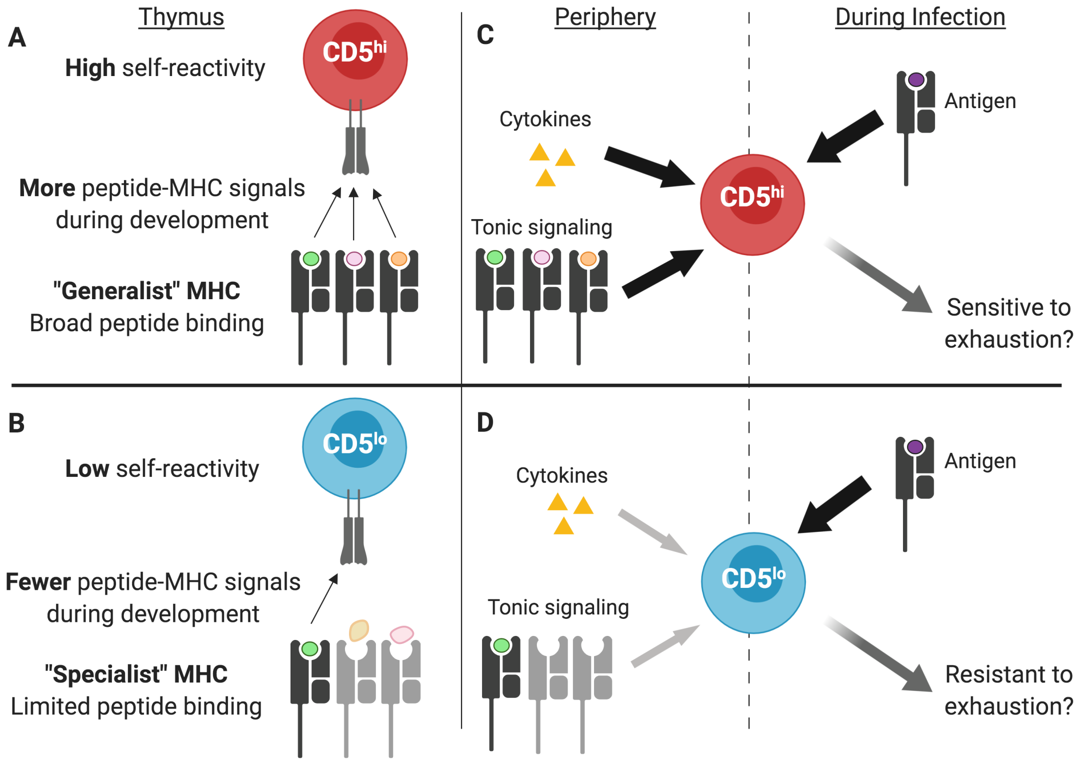


**Supplementary Figure 7:** Model for MHC-1 specialists and CD8+ elite control. **(A)** MHC-1 generalists bind a large set of self-peptides allowing for positive selection of T cells with relatively high self-reactivity ($CD5^{high}$). **(B)** MHC-1 specialists display low and unstable surface expression due to poor binding of self-peptides, resulting in the positive selection of T cells with low self-reactivity ($CD5^{low}$). **(C)** CD8+ T cells with high self-reactivity are more responsive to cytokines and may experience greater TCR tonic signaling in the periphery, which may promote exhaustion during chronic infection. **(D)** In contrast, CD8+ T cells with low self-reactivity are less responsive to inflammatory environments and may receive lower tonic signals, allowing them to persist without becoming exhausted during chronic infection. Figure created with BioRender.
